# Contrast gain control is a reparameterization of a population response curve

**DOI:** 10.1101/2024.07.29.605608

**Authors:** Elaine Tring, S. Amin Moosavi, Mario Dipoppa, Dario L. Ringach

## Abstract

Neurons in primary visual cortex (area V1) adapt in different degrees to the average contrast of the environment, suggesting that the representation of visual stimuli may interact with the state of cortical gain control in complex ways. To investigate this possibility, we measured and analyzed the responses of neural populations to visual stimuli as a function of contrast in different environments, each characterized by a unique distribution of contrast. Our findings reveal that, for a given stimulus, the population response can be described by a vector function **r**(*g*_*e*_*c*), where the gain *g*_*e*_ is a decreasing function of the mean contrast of the environment. Thus, gain control can be viewed as a reparameterization of a population response curve, which is invariant across environments. Different stimuli are mapped to distinct curves, all originating from a common origin, corresponding to a zero-contrast response. Altogether, our findings provide a straightforward, geometric interpretation of contrast gain control at the population level and show that changes in gain are well coordinated among members of a neural population.

## Introduction

Neurons in primary visual cortex respond to changes in the mean contrast of the environment by rigidly shifting their contrast-response function along the log-contrast axis (Ohzawa et al. 1982). The functional goal of such adaptation is to align the region of maximal sensitivity with the geometric mean of contrasts observed in the recent stimulus history and maximize information transmission (Laughlin 1981). To illustrate this effect, consider *r*(*c*) to be the contrast-response function of a neuron measured in an environment with a specified average contrast (**Fig 1A**, yellow curve, vertical arrow shows mean contrast). The contrast-response curve in a new environment with a higher, mean contrast would be shifted to the right (**Fig 1A**, orange curve). The transformed response can be described by *r*(*gc*), where *g* < 1 represents a reduction in contrast gain (Geisler and Albrecht 1992; Heeger 1992; Ohzawa et al. 1982).

**Figure 1.**
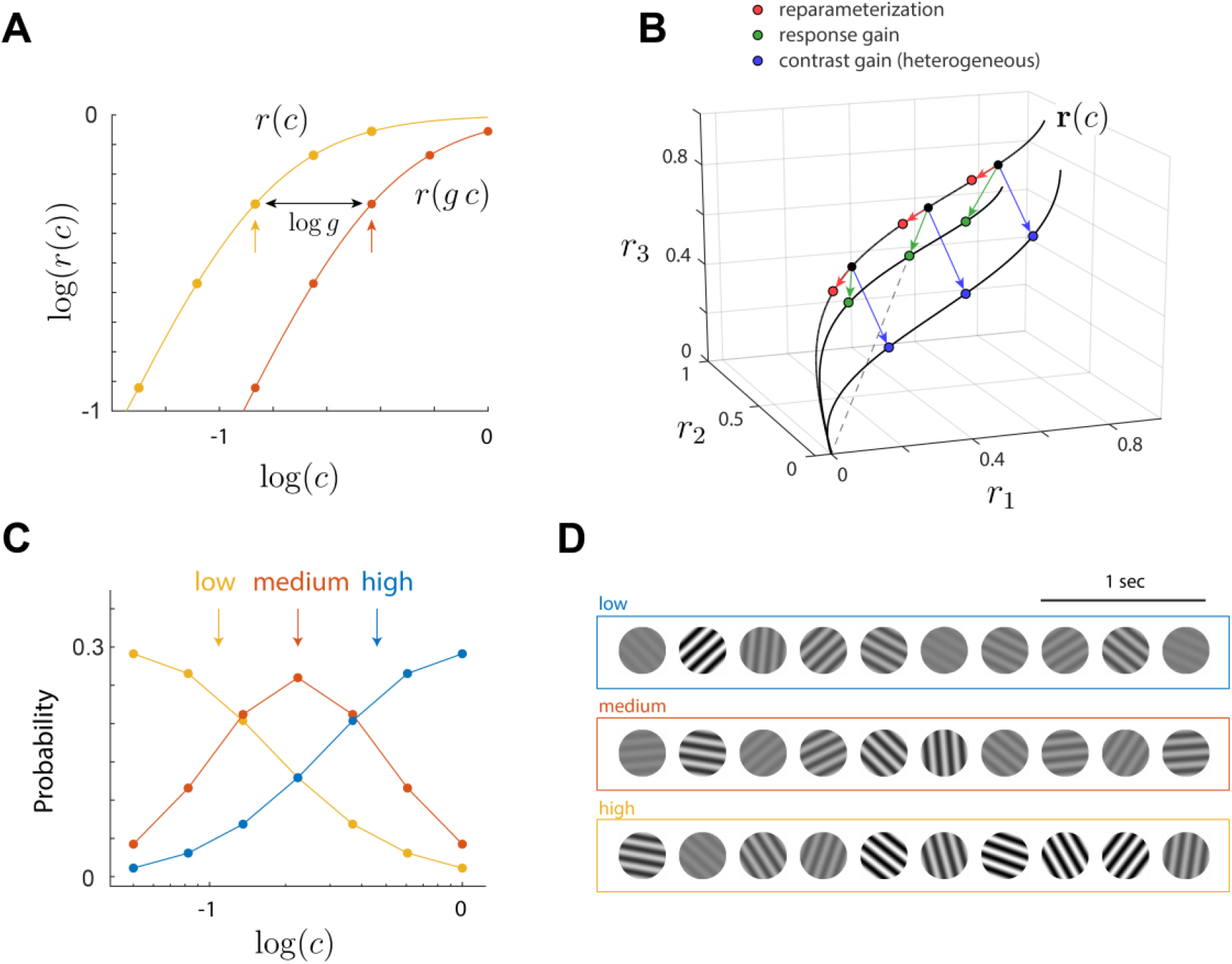
Gain control, hypothesis and experimental design. **A**. Early studies of contrast gain control measured the response of a single neuron in environments composed of five different contrast levels equally spaced along the log contrast axis (*Ohzawa et al. 1985*). Yellow and orange dots represent the responses obtained in two different environments, and the geometric mean within each environment is indicated by the vertical arrows. Here and in the sequel, logarithms of contrast values are expressed in base 10, with 0 anchored at 100%. The result of such an experiment is that the contrast response curves are shifted horizontally such that the geometric mean of the contrast aligns with the center of the curve. Thus, if *r*(*c*) expresses the responses in the yellow environment, then the responses in the orange environment with a higher mean contrast will be shifted to the right and can be described by *r*(*g c*). The horizontal shift between the curves is *log g*. Perfect adaptation occurs when *g* equals the ratio between the mean contrasts of the environments, but partial adaptation is also possible. **B**. Different scenarios for how gain control could act at the population level. The responses of three hypothetical neurons in one environment at three different contrast levels are depicted by solid black dots. The dots lie on a curve **r**(*c*). Reparameterization predicts that switching to an environment with higher, mean contrast, would result in the responses shifting towards the origin (red arrows/dots). Response gain posits that responses will scale down towards the origin (green arrows/dots). The dashed line shows the response will lie along the line joining the origin and the original population response. If neurons experience heterogenous changes in contrast gain the points ought to move to different contrast response curve (blue arrows/dots). **C**. Definition of environments. Our experimental design consists of three environments defined by distinct contrast distributions on a fixed set of contrast levels covering the range from 5 to 100%. The geometric means of the distributions (vertical arrows) differ due to the different frequency of presentation of each contrast level. The contrast levels are the same for all environments (as opposed to the design from earlier studies **A**). We used three distributions corresponding to high (blue), medium (orange) and low (yellow) mean contrast values. **D**. Examples of stimulus sequences for the different experimental conditions. Within each environment, the contrast of gratings is drawn according to their respective distributions in **C**, while their orientation and spatial phase are uniformly distributed. The sequences were presented at a rate of 3 stimuli per sec.

We hypothesize that such relationship generalizes to neural populations. Namely, we postulate the response of a population under contrast gain control is **r**(*g*_*e*_*c*), where **r** is a vector function, *c* is the stimulus contrast, and *g*_*e*_ is a gain factor that decreases monotonically with the mean contrast of the environment. Mathematically, this represents a linear reparameterization of a single contrast-response curve. It is helpful to depict the working of reparameterization graphically. Suppose the hypothesis holds and that we measure the responses of the population at three contrast levels (**Fig 1B**, black dots). These points will lie on the contrast response curve of the population under the current environment, **r**(*c*) (**Fig 1B**, black curve). Imagine we proceed to switch the environment to one with a higher mean contrast. Under reparameterization the responses of the population at the same contrast levels will shift along the curve towards the origin (**Fig 1B**, red arrows and dots). From a coding perspective, the advantage of reparameterization is that a visual stimulus generates a single response curve which remains invariant between environments. Assuming different stimuli generate different curves, the identification of stimulus by downstream areas can be reduced to the question of whether a given population response lies on a stimulus curve. Implementing reparameterization requires the neurons in the population to adjust their gains by the same factor. If neurons respond to a change in the environment by modifying their gains in very different amounts, the responses will move to move to a different response curve, entangling the representation of visual stimuli with the state of cortical gain control (**Fig 1B**, blue arrows and dots). In this scenario, the contrast response for a given stimulus depends on the state of gain control, complicating the decoding of visual information by downstream visual areas. Finally, it may be possible for changes in the mean contrast of the environment to induce modulation in response gain (Albrecht and Hamilton 1982; Ferguson and Cardin 2020; Hamilton et al. 1989; Sclar et al. 1989). In this case, the transformed response is given by *g*_*e*_**r**(*c*), with *g*_*e*_ representing, once again, a gain factor that decreases with the mean contrast of the environment. The signature of response gain is that while the magnitude of a population vector is changes between environments, its direction remains constant (**Fig 1B**, green arrows and dots). Such a coding strategy is appropriate to generate an invariant representation of stimulus contrast, as one can then identify the direction of population vectors with the absolute contrast of a stimulus independent of the environment.

At first glance, the reparameterization hypothesis appears to be on shaky grounds, as V1 neurons exhibit considerable diversity when studied individually – some cells show robust changes in gain as a function of mean contrast, while others do adapt at all (Ohzawa et al. 1982, 1985). However, in these studies, responses were measured independently at each neuron’s optimal orientation, spatial and temporal frequencies. The choice of stimulus parameters could have affected a normalization signal, such as average cortical activity, which in turn controls contrast gain (Carandini et al. 1997; Carandini and Heeger 2011; Heeger 1992; Schwartz and Simoncelli 2001). Thus, using stimulus parameters that are average for the population could generate a larger normalization signal than using extreme values. Thus, the diversity of stimulus parameters could be partly responsible for the mixture of gain changes observed in prior, single cell data. Instead, to address the reparameterization hypothesis directly and to circumvent problems in the interpretation of past data, we set out to measure and analyze the contrast response of a neural population to a fixed visual stimulus under different experimental conditions or “environments”, each associated with a unique distribution of contrast values with different means (**Fig 1C,D**).

To anticipate the results, we find that the population response to a visual stimulus is well captured by a single, vector function **r**(*g*_*e*_*c*). Moreover, we find that 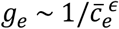 (with *ϵ* > 0), where 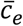 is the geometric mean of the contrast in the visual environment. Thus, gain control at the population level admits a simple description as a linear reparameterization of a contrast-response curve. A visual pattern can be identified with its unique contrast-response curve **r**(·), which is invariant across environments, thereby facilitating downstream decoding. Different visual patterns generate distinct curves, all originating from a common origin representing a zero-contrast response. These findings indicate that contrast gain must be reasonably coordinated across all cells in a cortical population. Our results offer a simple, geometric interpretation of contrast gain control at the level of neural populations.

## Materials and Methods

### Experimental Model and Subject Details

All procedures were approved by the University of California, Los Angeles (UCLA)’s Office of Animal Research Oversight (the Institutional Animal Care and Use Committee). The experiments also complied with the guidelines set by the U.S. National Institutes of Health on animal research. A total of 9 mice, male (4) and female (5), aged P35-56, were used. These animals were obtained as a cross between TRE-GCaMP6s line G6s2 (Jackson Laboratory, https://www.jax.org/strain/024742) and CaMKII-tTA (https://www.jax.org/strain/007004). There were no obvious differences in the results between male and female mice.

### Surgery

We measured cortical activity using two-photon imaging through cranial windows implanted over V1. Carprofen was administered pre-operatively (5 mg/kg, 0.2 mL after 1:100 dilution). Mice were anesthetized with isoflurane (4–5% induction; 1.5–2% surgery). Core body temperature was maintained at 37.5°C. We coated the eyes with a thin layer of ophthalmic ointment during the surgery to protect the corneas. Anesthetized mice were mounted in a stereotaxic apparatus using blunt ear bars placed in the external auditory meatus. A section of the scalp overlying the two hemispheres of the cortex was then removed to expose the skull. The skull was dried and covered by a thin layer of Vetbond and an aluminum bracket affixed with dental acrylic. The margins were sealed with Vetbond and dental acrylic to prevent infections. A high-speed dental drill was used to perform a craniotomy over monocular V1 on the left hemisphere. Special care was used to ensure that the dura was not damaged during the procedure. Once the skull was removed, a sterile 3 mm diameter cover glass was placed on the exposed dura and sealed to the surrounding skull with Vetbond. The remainder of the exposed skull and the margins of the cover glass were sealed with dental acrylic. Mice were allowed to recover on a heating pad and once awake they were transferred back to their home cage. Carprofen was administered post-operatively for 72 h. We allowed mice to recover for at least 6 days before the first imaging session.

### Two-photon imaging

Imaging sessions took place 6–8 days after surgery. Procedures were identical to those described earlier (Tring et al. 2024). Mice were positioned on a running wheel and head-restrained under a resonant, two-photon microscope (Neurolabware, Los Angeles, CA). The microscope was controlled by Scanbox acquisition software and electronics (Scanbox, Los Angeles, CA). The light source was a 920 nm excitation beam from a Coherent Chameleon Ultra II laser (Coherent Inc., Santa Clara, CA). We used a x16 water immersion objective for all experiments (Nikon, 0.8 NA, 3 mm working distance). The microscope frame rate was 15.6 Hz (512 lines with a resonant mirror at 8 kHz). The field of view was 730 µm × 445 µm in all sessions. The objective was tilted to be approximately normal on the cortical surface. Images were processed using a pipeline consisting of image registration, cell segmentation, and signal extraction using Suite2p (Pachitariu et al. 2017). A custom deconvolution algorithm consisting of linear filtering followed by half-rectification and a power function was used (Berens et al. 2018).

### Visual stimulation

A Samsung CHG90 monitor, positioned 30 cm in front of the animal, was used for visual stimulation. The screen was calibrated using a Spectrascan PR-655 spectro-radiometer (Jadak, Syracuse, NY), generating gamma corrections for the red, green, and blue components via a GeForce RTX 2080 Ti graphics card. Visual stimuli were generated by a custom-written Processing 4 sketch using OpenGL shaders (see http://processing.org). At the beginning of each experiment, we obtained a coarse retinotopy map of the cortical section under study (Tring et al. 2022). The center of the aggregate population receptive field was used to center the location of our stimuli in these experiments. Stimuli were presented within a circular window with a radius of 25°.

We used a sequence of flashed, sinusoidal gratings presented at a rate of 3 per second for stimulation. The spatial frequency was fixed at 0.04 cycles per deg, matching the average of the V1 population (Niell and Stryker 2008). Sequences were presented in blocks representing one among three possible environments with different contrast distributions (**Fig 1C**). The distributions were truncated log-normal sampled on a discrete set of seven contrast levels 5% × *ξ*^*q*^ for *q* = 0,1, …, 6 and *ξ* = 1.6475. Thus, contrast levels were equally spaced in logarithmic steps. The geometric means of the contrast in the three environments were 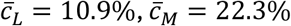, and 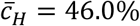 (**Fig 1C**, vertical lines), which we refer to as the low (*L*), medium (*M*) and high (*H*) contrast environments. Thus, the average contrasts of the environments are spaced by an octave. Note that while the mean contrasts in the environments differ, their range is the same, as opposed to the experimental design used in earlier studies (**Fig 1A**). This allows us to compute the responses over the entire range of contrasts in all environments. Stimulus sequences were generated by uniformly drawing the orientation and spatial phase of the grating, while drawing the contrast from the corresponding environment distribution (**Fig 1D**). All six permutations of {*L, M, H*} − environments were presented in a randomized order, leading to a total of 18 experimental blocks. Each block was presented for 5 min, for a total of 900 stimuli per block. Each environment appeared 6 times during the session, resulting in 5400 stimuli per environment. A one-minute blank screen was presented between blocks. The presentation of each grating was signaled by a TTL pulse sampled by the microscope. As a precaution, we also signaled the onset of the stimulus by flickering a small square at the corner of the screen. The signal of a photodiode at that location was sampled by the microscope as well.

### Definition of population responses

For each environment *e* ∈ {*L, M, H*}, and orientation *θ*, we calculated the mean response, **r**_*e*_(*θ, c, T*), averaged over spatial phase, *T* microscope frames after the onset of the stimulus. Here, *e* ∈ {*L, M, H*} is one of the environments (**Fig 1B**), *θ* represents the orientation of the grating, and *c* its contrast. The response averaged across all orientations is denoted by **r**_*e*_(*c, T*). As we will see, the largest response magnitude is obtained for *e* = *L* and *c* = 100%. Thus, we define the optimal time-to-peak, *T*_*opt*_, as the one for which the Euclidean norm of **r**_*L*_(100%, *T*) attained its peak after stimulus onset. Across all our sessions we found *T*_*opt*_ = 287 ± 33 msec (mean ± 1SD, *n* = 17). We define **r**_*e*_(*θ, c*) = **r**_*e*_(*θ, c, T*_*opt*_) and similarly, **r**_*e*_(*c*)= **r**_*e*_(*c, T*_*opt*_). The Euclidean norm of these vectors will be denoted by *r*_*e*_(*θ, c*) and *r*_*e*_(*c*) respectively.

There is a technical point concerning the estimate of population norms that deserves attention. We want to estimate the Euclidean norm *r* of the mean population response to *m* trials of a stimulus in a population of *d* independent neurons. The response of the neurons in any one trial is the realization of a random variable *r*_*i*_, with *i* = 1, …, *d*. The actual response of neuron *i* to the stimulus in trial *k* will be denoted by 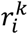. If we let 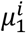 represents the mean response of the *i* − *th* neuron, we want to find an estimate of 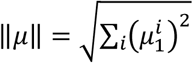 from the data 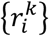.

A reasonable way to proceed is to estimate the squared norm of the population response as 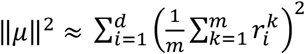. However, it is easy to see this estimate is biased. First, let us consider the inner term, which expands to 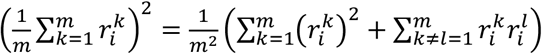. What would be the average value of this quantity if we were to repeat the experiment many times? If we take expected values on both sides of the equation we obtain 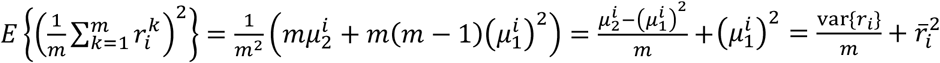. Here, 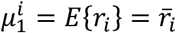 represent the mean of *r*_*i*_, and 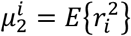 is its second moment. Thus, on average, we there is a bias term 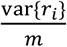 which depends on the number of trials. To correct for it, we calculate the sample variance and subtract the term 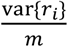 from 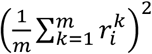. Finally, to obtain an estimate of the norm, we add all the terms across neurons and take the square root. Such bias correction was applied to the calculation of *r*_*e*_(*θ, c*).

### Bézier curve fit to contrast response data

For a fixed visual pattern, we obtain the mean response to 7 different contrast levels in 3 different environments (a total of 21 points). As we show in **Results**, these points can be projected into the first two principal components without major geometric distortions. These data appear to lie on a single curve. To characterize the shape of such a curve we fit a quadratic Bézier curve as follows. First, the data are normalized so that the response with minimum norm is mapped to (0,0), while the response with the largest norm is mapped to (0,1). We achieve this with a similarity transformation, which does not distort the shape of the curve. A quadratic Bézier curve has 3 control points. We choose the first one to align with the origin, **p**_0_ = (0,0), and the third one to be **p**_2_ = (0,1). The only free parameter left is the point **p**_1_, which leads to the Bézier curve: ***B***(*t*) = *2t*(1 − *t*)**p**_1_ + *t*^2^(0,1). For a given choice of **p**_1_ we can compute the minimum distance to the curve from each data point. For a given choice of **p**_1_, we define the error of the fit as the average mean square distance from the data points to the curve. Then, we minimize the error as a function of **p**_1_ using Matlab’s fminsearch. There is nothing special about the use of Bézier curves, other interpolation methods could have worked as well to illustrate that the responses lie along a smooth curve (**Fig 3**).

### Rigor and reproducibility

We conducted experiments by independently measuring the adaptation of V1 populations in *n* = 17 independent sessions. Linear models were fitted to the data using Matlab’s `fitlm` function. The goodness of fit of linear models was evaluated using the coefficient of determination, *R*^2^. As the study did not involve different groups undergoing different treatments, there was no need for randomization or blind assessment of outcomes. Data selection was used to process all putative neurons selected by Suite2p (Pachitariu et al. 2017) and select only those that responded significantly to visual stimulation. This was done by calculating the ratio between the response of neurons at the optimal time for the population, *T*_*opt*_, and their baseline response just prior to the onset of stimulation. We selected neurons for which such ratio was larger than 8. No selection was done with respect to the tuning of neurons for orientation – both cells with good and poor orientation selectivity were included. The median number of cells in our populations was 110, with the first and third quartiles at 210 and 335 respectively.

## Results

We begin by providing low dimensional visualizations of the geometry of **r**_*e*_(*θ, c*) using principal component analysis. These analyses expose some potential features of gain control, including the reparameterization of a single contrast-response curve. As low-dimensional visualizations can incur geometric distortions, subsequent analyses are performed in native response space. In this context, we investigate the structure of the pairwise Euclidean distance matrix 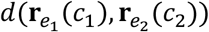 between responses across environments. To test the reparameterization hypothesis, we show that given two different environments *e*_1_ and *e*_2_, where the geometric mean of contrast in *e*_2_ is lower than *e*_1_, we can find a constant *g* < 1 such that 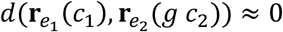. This finding implies that 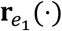 and 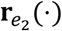 lie approximately on a single response curve, the hallmark of reparameterization. We then provide a statistical model for the dependence of the response magnitude *r*_*e*_(*c*) with changes in the environment, showing that 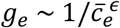, where 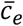 is the geometric mean of the contrast in the visual environment. We then consider and reject the response gain model as a good contender to explain the data. Finally, we relate the properties of the population to those of single neurons and provide an estimate of the degree of coordination of gain control in the population in response to a fixed visual stimulus.

### Low dimensional visualization of population responses

To visualize the structure in our datasets we first used principal component analysis to examine the structure of **r**_*e*_(*θ, c*) in three dimensions (**Fig 2A**). Different colors are used to show data from different orientations. Each environment is assigned a different symbol (low = circles, medium = asterisks, high = squares). For a fixed environment and orientation, straight lines join data points at adjacent contrast levels. The resulting curves show the shape of **r**_*e*_(*θ, c*) as a function of *c* for the different environments and stimulus orientations.

**Figure 2.**
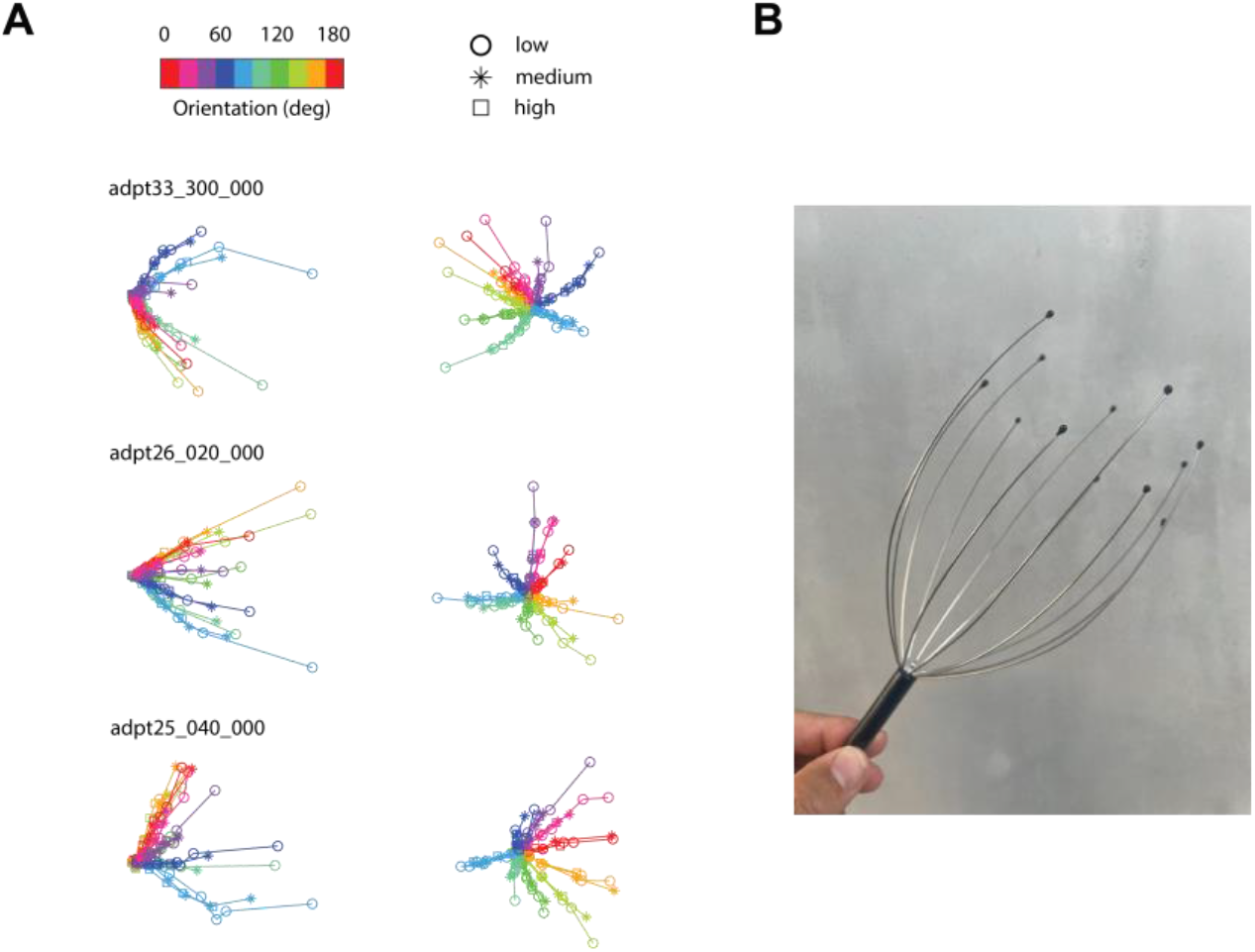
Low dimensional visualization of population responses. **A**. Each row corresponds to data from a different experimental session, showing two views of the projection of the data into the first three principal components. Each color corresponds gratings of different orientations (top color bar), while each symbol represents data for environments with different mean contrasts (circles = low, asterisk, medium, square = high). The contrast response for a fixed orientation across different orientations all lie approximately on the same curve. Mean responses for low contrasts converge into a point. **B**. A visual that captures the overall structure of the data. Each “arm” of the scalp massager represents a contrast response curve for a fixed orientation. Changing environments causes a shift in the mean responses along the arm, which implies gain control is a reparameterization of a single curve.

Some salient features in the structure of the data catch the eye. First, for any given orientation, the contrast response functions across different environments overlap substantially – all symbols representing the data for a single orientation at different contrast levels and environments appear to lie approximately on a single curve. This is consistent with the idea that gain control acts to reparametrize the population response (**Fig 1B**). In other words, changing the environment only shifts the data along the curve and does not move them “off the manifold” associated with a given orientation (Jazayeri and Afraz 2017). Second, different orientations generate different curves emanating from a common origin. This type of structure is expected from the orientation tuning of cortical neurons and the fact that as contrast is decreased, we expect the responses at all orientations to converge to the response to a zero contrast “origin”. Third, the contrast response curves resemble straight rays at low contrast values but show clear curvature at moderate to high contrast values. The curvature is mostly visible in data from the low-contrast environment, which also generates the responses with the largest magnitudes. One can think of the geometry of the responses as resembling a scalp massager, with each wire representing the “arms” generated by stimuli at different orientations (**Fig 2B**). The arms remain invariant with changes of the environment. Of course, these observations should be interpreted with care, as the fraction of variance captured by a projection into the first three principal components is only about half of the total (0.54 ± 0.056, mean ± 1SD).

### Contrast responses for a given stimulus across different environments lie on a curve

Our visualization of the data by principal component analysis (PCA) in 3D is likely to be dominated by the need to capture the disparate responses evoked by different orientations. Instead, to focus on the analysis of the shape of the contrast response functions, we performed PCA on the individual “arms” of the dataset, each representing the responses in all 3 environments for 7 contrast levels at a single orientation. Our analyses show that projecting the data into the first two principal components accounts for 0.83 ± 0.033 (mean ± 1SD) of the total variance of the arms, reasonably capturing their shape (**Fig 3**). Thus, contrast response curves lie mostly on a plane. The projected data points in 2D were normalized using a similarity transformation, such that the response with the smallest norm was mapped to (0,0) and the one with largest norm mapped to (0,1). This was done to allow the comparison of the shape of contrast response functions for different orientations.

**Figure 3.**
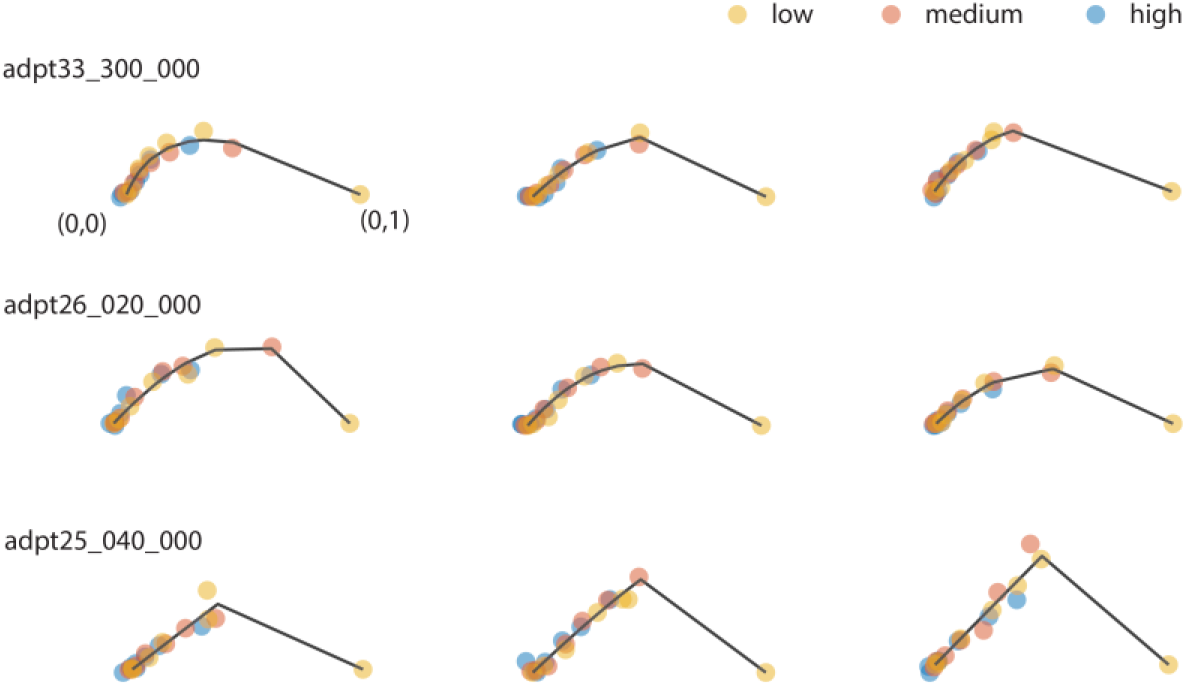
Contrast response to a fixed visual pattern across different environments lie approximately on a 2D curve. Each row shows the structure of some “arms” in one experimental session. For each arm, the data points represent the projection of the mean responses at a fixed orientation for varying levels of contrast for the three environments (low, medium and high contrast) onto the first two principal components. Each environment is coded by a different color. Only three orientations per session are shown. Each arm is normalized so the response with minimum norm is mapped to (0,0) and the one with largest response to (0,1). Dark segments join the locations on the Bezier curve fit that are closest to the data.

A key observation is that data points for each arm tend to fall along a single curve (**Fig 3**). The largest responses (farther from the origin) are obtained for the highest contrast level in the low-contrast environment. Switching from the low contrast environment to a medium contrast environment merely shifts the points along the curve towards the origin at (0,0). The same occurs as we move from the medium contrast environment to the high contrast environment. This finding is consistent with the hypothesis that gain control serves to reparametrize the population responses. Moreover, as we will see below, this reparameterization has a simple dependence on the mean contrast of the environment. We note that in some experiments, the data lied almost entirely along a ray except for the point at the highest contrast in the low-contrast environment (**Fig 3**, bottom row). This likely resulted from our experimental design under-sampling the 60-100% contrast range, a limitation we plan to correct in future studies.

### Testing reparameterization using distance matrices in native space

Next, to circumvent any possible distortions in the structure of the data incurred by dimensionality reduction methods, we test the reparameterization hypothesis directly in native space. By “native space” we mean the Euclidean, *d* dimensional space, where the response vectors live. We achieve this by studying the structure of pairwise distance matrices 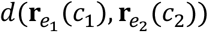 as follows. Let us assume, without loss of generality, that the mean of contrast in *e*_2_ is lower than *e*_1_. Thus, responses in *e*_2_ at any one contrast are larger than the ones obtained in *e*_1_. The linear reparameterization hypothesis predicts that the response to *c*_1_ in *e*_1_ can be matched by reducing the contrast in *e*_2_ by a fixed gain factor, 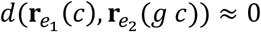. Indeed, when plotting *d*(**r**_*med*_(*c*_1_), **r**_*low*_(*c*_2_)) and *d*(**r**_*high*_(*c*_1_), **r**_*med*_(*c*_2_)) we observe that minimum distances fall approximately on a diagonal parallel and displaced from the identity line. As contrast axes are logarithmic, the displacement simply represents a multiplicative effect of the environment on contrast – the signature of gain control. The same effect seen in the structure of *d*(**r**_*high*_(*c*_1_), **r**_*low*_(*c*_2_)). Here, the shift of the diagonal doubles, matching a doubling in the ratio between the mean contrasts of the environments. Altogether, we conclude that for any two environments we can find a factor *g* such that *d*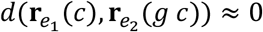. One caveat is that both measurement noise and the coarse sampling of contrast values provide a lower bound on how close to zero this value can get. In these analyses, distances were normalized by the diameter of the dataset (the maximum distance between any two responses), and the minimum distances obtained were 0.096 ± 0.022 (mean ± 1SD).

### Dependence of gain on the mean contrast of the environment

How does the gain *g* change as a function the mean contrast of the environment? Although the location of the diagonals along which the distance is minimum offers one approach (**Fig 4**), it is limited by the coarse sampling of contrast in the data. Here, we offer an approach that relies on the comparison of response magnitudes between environments. Note that if 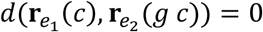, then it must be the case that the magnitudes satisfy 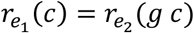. Therefore, in a way analogous to the study of single neurons, we should see a horizontal shift in the *magnitude* of population responses between environments by *log g* (**Fig 1A**). Indeed, when plotting the population magnitudes as a function of contrast in double logarithmic axis they appear as shifted lines (**Fig 5**). This analysis also replicates our previous finding that the magnitudes of population responses are a power law of contrast, *r*(*c*) *∼ c*^*δ*^ (Tring et al. 2024). Unfortunately, the shift in the population norm is ambiguous, as it could potentially be interpreted as either a horizontal or vertical shift of a line, corresponding to changes in contrast gain, response gain, or a mixture of both (Albrecht et al. 1984a, 1984b; Hamilton et al. 1989). However, as we argue below, a horizontal shift is the one consistent with the structure of the data matrix and the geometry of the responses. For the moment, let us describe the transformation observed as changes in contrast gain. Note that magnitudes of the shifts are approximately equal as we move from low-contrast to medium-contrast, and from medium-contrast to high-contrast environments, representing equal steps in log-contrast (**Fig 1C**). Thus, it is natural to put forward a gain control model 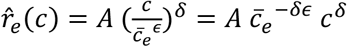, where 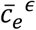is the geometric mean of the contrast in *e*. As it turns out, this model performs extremely well (**Fig 4**, *R*^2^ = 0.976 ± 0.01, mean ±1SD, *n* = 17). Across all sessions, we obtain *δ* = 0.81 ± 0.092 and *ϵ* = 0.68 ± 0.06 (mean ±1SD). Altogether, we conclude the response of the population to a fixed visual pattern in environment *e* is given by **r**(*g*_*e*_*c*) with 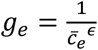.

**Figure 4.**
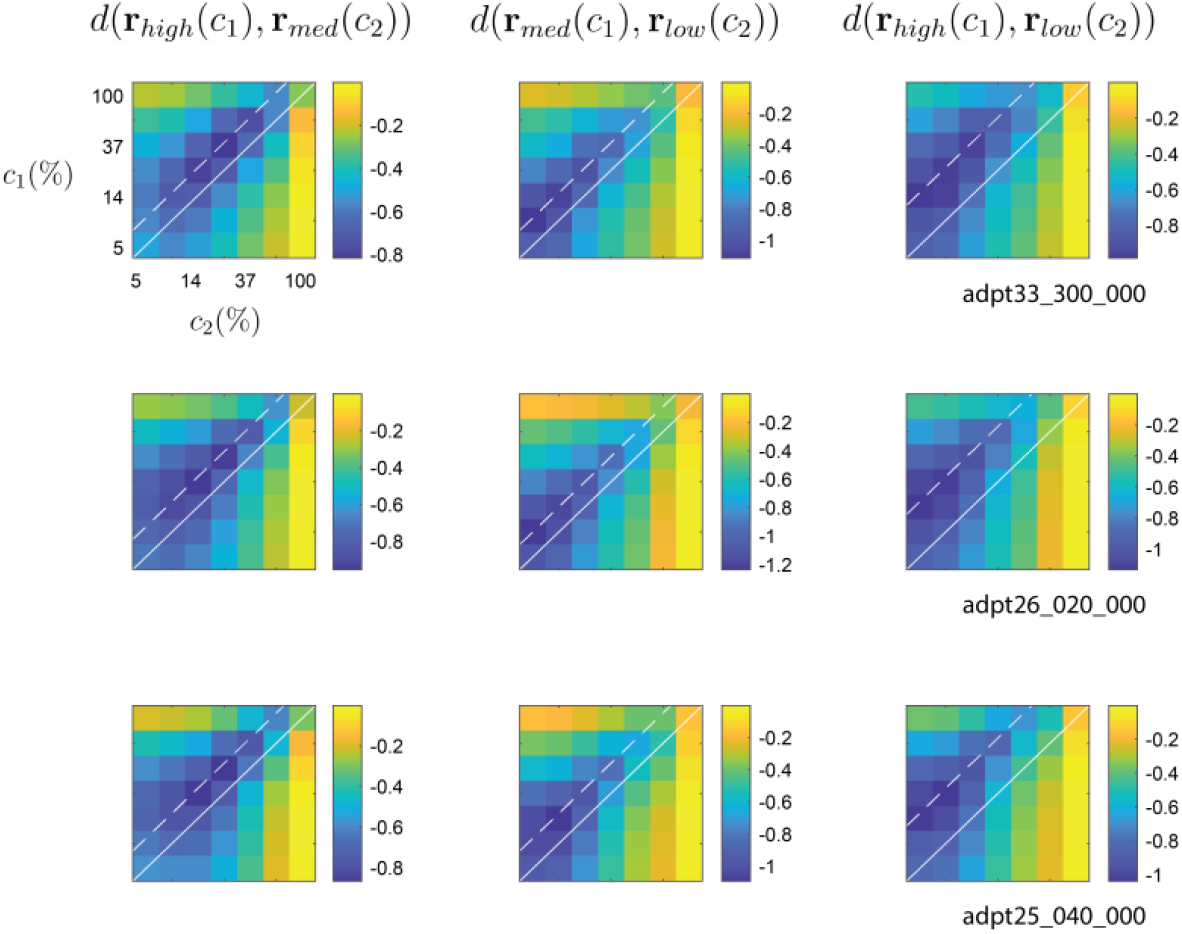
The structure of distance matrices is consistent with reparameterization. Each row, representing data from different sessions, shows the Euclidean distance between population responses across different environments. The first column shows the distance between responses in high and medium contrast environments; the second column between medium and low environments; and the third column between high and low environments. Distances are normalized to the diameter of the dataset and displayed on a logarithm (base 10) scale. We observe that distance minima lie on a diagonal parallel to the identity line. The dashed line represents the location of the diagonal on which the average distances reach a minimum. Solid white line is the identity line. Such diagonal structure is consistent with the linear reparameterization hypothesis.

**Figure 5.**
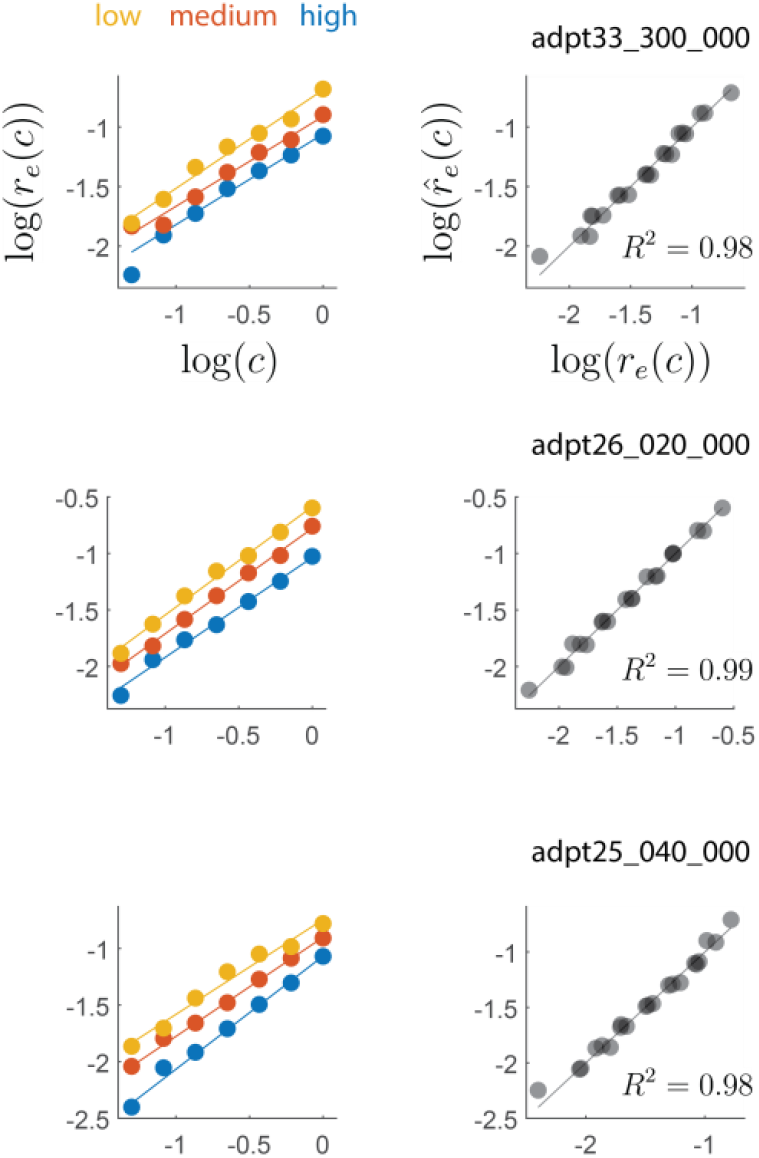
Dependence of gain on the mean contrast of the environment. Each row represents data from sessions matching those in prior figures. The right column shows the magnitude of the responses as a function of contrast for the different environments. The plot is in double logarithmic axes. The solid lines represent the best linear fits to the data from each environment (fit independently for each environment). The lines have approximately the same slope and are shifted in about equal amounts. This suggests that the entire dataset may be captured by the model 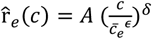. The panels on the right show the fit of such a model to the data. The quality of the fits is very good, with *R*^2^values ∼0.98. Solid gray line represents the identity line.

### Changes in response gain are inconsistent with the geometry of the data

Let us now go back and discuss the ambiguity between changes in response gain versus changes in contrast gain in the response magnitude data (**Fig 5**). The response gain model is represented by the relationship *g*_*e*_ **r**(*c*), where the output gain is modulated by the environment (**Fig 1B**). This implies that, for any given contrast, the *direction* of population response should not change across environments – only its magnitude does. Thus, one would predict that the cosine distance matrix 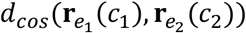 should be zero along its diagonal, when *c*_1_ = *c*_2_. Instead, we observe the directions of the population vectors across two environments are most similar at different contrast values (**Fig 6**), in parallel to behavior of the Euclidean distance matrix (**Fig 4**). Thus, we safely conclude the data does not conform to the response gain model.

**Figure 6.**
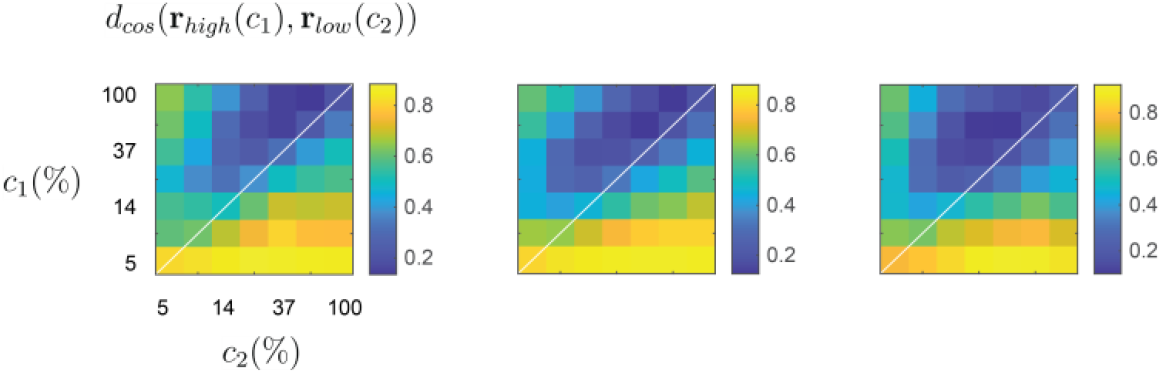
Changes in response gain are inconsistent with the geometry of the data. Each panel shows the cosine distance between responses in the high and low environments in independent sessions. While response gain predicts distances should be near zero along the main diagonal, we observe the minima off the main diagonal in parallel with the behavior of the Euclidean distance matrices (**Fig 4**).

### Assessment of the dispersion of gain control in a population

In principle, reparameterization strictly holds only when the changes in gain in the population are identical. Of course, some degree of variability is expected in the data. Here, we assess the degree of gain control dispersion across neurons for a fixed visual stimulus across environments and how it relates to the analyses at the population level.

For any given session, we analyzed data obtained at a single orientation – one of the “arms”. In each arm, only a small fraction of cells will have a preferred orientation matching that of the stimulus and respond vigorously. Many other neurons respond with weak and noisy responses. If we analyze the contrast response functions of the responsive neurons, we see that some recapitulate, at least partially, what is observed at the population level (**Fig 7A**). One common departure is that responses saturate at the highest contrast levels for the low-contrast environment (**Fig 7A**, top row, yellow data points). Nonetheless, the responses are reasonably fit by the model 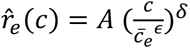, which allows us to estimate the shift in the contrast response function of each neuron across environments (which equals *ϵ log* 2, as the mean contrast between the environments differ by one octave). We can quantify the dispersion in the shifts by the coefficient of variation (CV) of *ϵ*. When pooling data across different arms and different experimental sessions, we find that, on average, *CV* = 0.27 ± 0.10 (mean ± 1SD, computed over 53 different “arms”) (**Fig 7B**).

**Figure 7.**
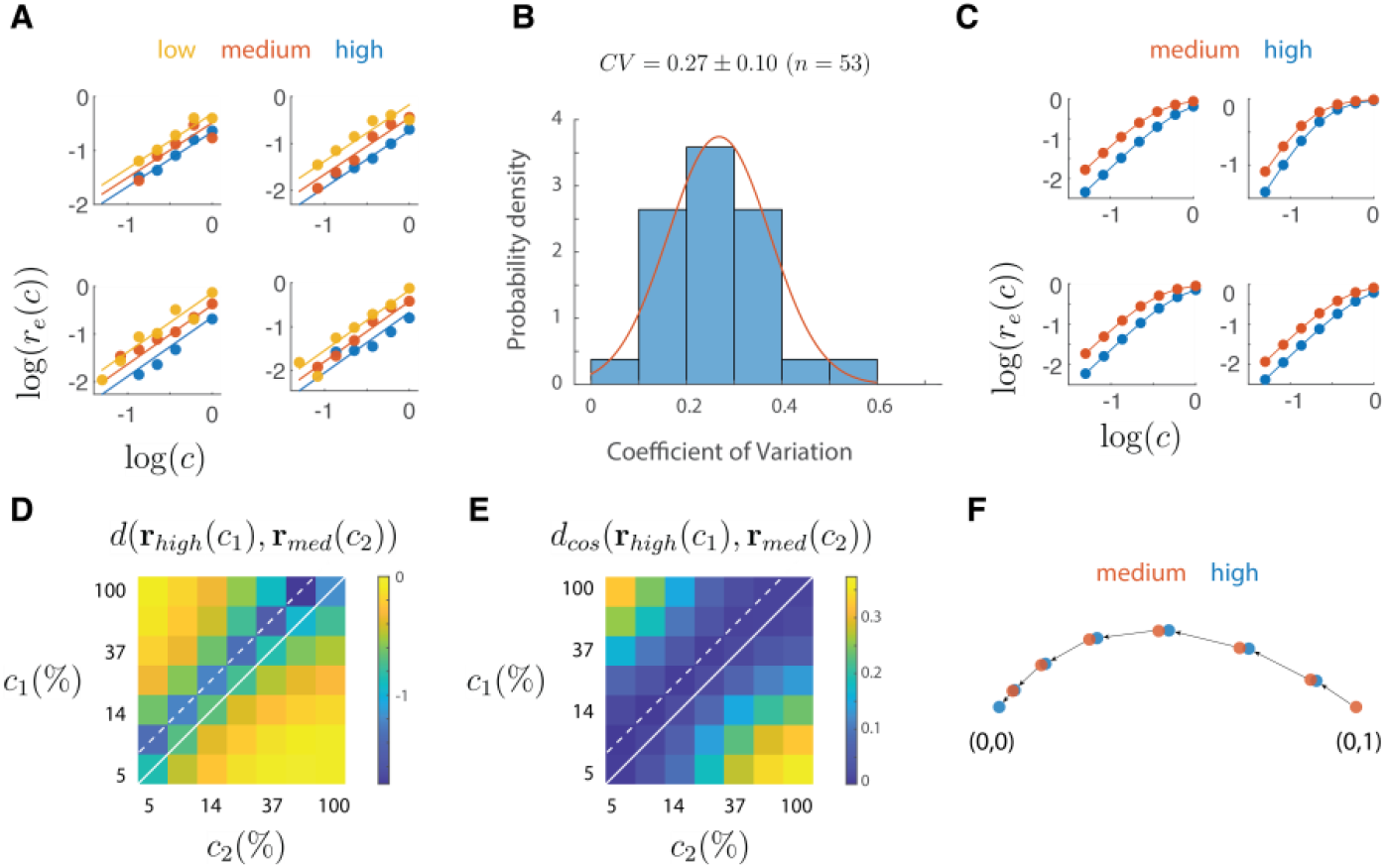
Observed dispersion in gain control across V1 neurons is consistent with reparameterization. **A**. Example of contrast response functions of single neurons for a fixed orientation. The solid lines are the fits of the model. The horizontal shift between the responses in different environments yields an estimate of the change in gain control. To assess the dispersion of gain control we compute the coefficient of variation of the shifts in a population of neurons. **B**. Distribution of the coefficient of variation pooled across different “arms” and imaging sessions. Only arms which included at least three responsive neurons in an arm were used. The coefficient of variation was corrected for bias given the small numbers involved, such that 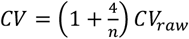, where *n* is the number of data points (Sokal and Rohlf 1995). Red curve represents a Gaussian fit. **C**. Simulated neurons with Naka-Rushton response profile and a gain shift distributed to match the mean coefficient of variation in **B**. The structure of the Euclidean (**D**) and cosine (**E**) distance matrices recapitulates the findings observed in the population analysis, with the minimum distances lying along a diagonal displaced from the unity line. **F**. The first two principal components capture 99% of the variance in the data, has a similar shape to those observed experimentally, and illustrates that a change in the mean contrast of the environment causes responses to shift along a single curve, consistent with reparameterization (**Fig 1**).

Is such a degree of variability small enough for the population to approximate reparameterization? We know the answer must be affirmative, as this is what the population analysis shows. Nonetheless, we can further corroborate this expectation with a simple exercise. We simulate a population of neurons in one environment by assuming the contrast response of the *i* − *th* neuron is given by the Naka-Rushton equation 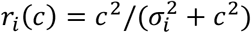 (Albrecht and Hamilton 1982; Naka and Rushton 1966). We selected *σ*_*i*_ to be distributed as a Beta function with *a* = 3 and *b* = 6, to match approximately the distribution seen in V1 (Sclar et al. 1990). We model a switch to a new environment of higher mean contrast by reducing the semi-saturation constants by a factor drawn from a triangular distribution designed to match the coefficient of variation of the data (**Fig 7C**). Now that we have contrast-response functions in the two environments, we can generate synthetic data and perform the same analyses as before, studying the structure of the pairwise Euclidean distance matrix (**Fig 7D**), the structure of the pairwise cosine distance matrix (**Fig 7E**), and generating a low-dimensional visualization of the transformation of responses resulting from a switch of environments (**Fig 7F**). We find that, for the experimentally observed degree of dispersion in gain control, the model reproduces the main phenomena seen in our data, including the shift of responses along a single curve which we accepted as is the signature of reparameterization (**Fig 7F**). Thus, a moderate level of coordination between neurons is sufficient to generate a representation that approximates reparameterization of responses by a population.

## Discussion

The central aim of this study was to test the hypothesis that contrast gain control can be interpreted as a linear reparameterization of the contrast-response function of the population (**Fig 1B**). Altogether, our data provide good support for this idea. First, a low dimensional visualization of the population responses indicated that responses at a fixed orientation as a function of contrast lie along a single “arm” when switching between environments (**Fig 2**). Second, analyzing the data separately for each arm showed that the response curves can be embedded in a plane and that the responses to different contrast levels shift along a single curve as the population as we modify the mean contrast of the environment (**Fig 3**). As these findings rely on analyses performed after dimensionality reduction, we sought to reveal signatures of reparameterization in the native response space as well. In this context, we demonstrated that the responses in one environment can be matched by scaling the contrast in a second one, as shown by minima of pairwise distance matrices lying along a diagonal displaced from the identity line (**Fig 4**). Finally, the data were well fit by population contrast-response function that is a power law of contrast (Tring et al. 2024), where contrast gain is a power law of the geometric mean of the environment (**Fig 5**). Such power law behavior emerges naturally in a population where with a wide distribution of semi-saturation constants. Altogether, our findings show that population responses under gain control can be viewed as the linear reparameterization **r**(*g_e_c*), with 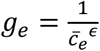.

Reparameterization simplifies the job of downstream areas, as the identity of a visual pattern is represented by its contrast response curve, which is invariant to changes in the distribution of contrasts in the environment. Finally, we note that perfect adaptation occurs for *ϵ* = 1. Our measured estimate of *ϵ* = 0.68 ± 0.06 means that, under the condition of our experiments, adaptation is only attained partially. Thus, coding of contrast is neither absolute (*ϵ* = 0) nor purely relative to the mean (*ϵ* = 1). Values of *ϵ* closer to one may be possible if we restrict contrast distributions in the experiments to those observed naturally (Mante et al. 2005).

In retrospect, a limitation of the study was the use of 7 log-steps from 5 to 100% to sample contrast. This choice left the range of contrast from 60 to 100% under sampled, which turned out to be the location the contrast response function appears to have the highest curvature (**Fig 2**). Taking a more detailed look at the shape of the contrast response curve calls for a denser sampling of contrast values, which will be remedied in future studies. Our findings are also limited to the truncated, log-normal distributions of contrast employed in these experiments (**Fig 1C**). It will be important to extend the range of environments by using distributions found in natural scenes to verify the behavior can be generalized (Clatworthy et al. 2003; Mante et al. 2005), including conditions where both contrast and mean luminance change (Geisler et al. 2007; Mante et al. 2005). In addition, in the present experiments we used a large, circular window for stimulation, which covered all the receptive fields of the population. Investigating the dependence of contrast gain with the spatial distribution of contrasts in the image is another important step in future studies (Albrecht et al. 1984a; Brady and Field 2000; DeAngelis et al. 1992; Schwartz and Simoncelli 2001). Such data may help link reparameterization to illusions of perceived contrast, such as the effect observed in the simultaneous contrast illusion (Carandini and Heeger 2011; Watson and Solomon 1997). Finally, some of the details of the geometry we recovered may be distorted by non-linear relationship between the actual spiking of neurons and our indirect inference from calcium imaging (Nauhaus et al. 2012). High-density electrophysiology will need to be conducted to assess any possible departures from imaging data.

We have not yet delved into the neural mechanisms implementing reparameterization, but it would have to address how the network can generate a reasonably coordinated change of gain in a local population. One prominent candidate is using a pooled cortical signal that controls gain in the local population (Heeger 1992), rather than by independent, self-calibration (Ullman et al. 1997). The ability to influence a pooled signal could also serve as a central “knob” that accounts for a separable power law relationship between the magnitude of the response with the probability of stimulus (Tring et al. 2023), its contrast (Tring et al. 2024) and, as shown in the present study, the mean contrast of the environment. Each of these factors appears to tweak the same gain “knob”, as demonstrated by the fact that a change in one can be compensated for by a change in another. For example, an increase in the probability of a stimulus can be counteracted by an increase in its contrast to keep the response magnitude constant (Tring et al. 2024).

We close by noting that the geometric view of contrast gain control is nothing more than a convenient way to look at the average behavior of a large population of neurons as they adapt to the contrast of an environment. The behavior of the population simply reflects the collective behavior of individual neurons. Other than a central mechanism that coordinates gain changes among neurons, there is no other “emergent” phenomena in the data. The advantage of our analyses is that it reveals lawful statistical relationships that describe the population responses accurately, even though many individual neurons respond weakly and generate noisy responses. An apt analogy would be a description of the behavior of gases in terms of statistical quantities such as pressure, temperature and volume. Here, we can obtain simple relationships between these quantities even though the behavior of the individual molecules can show considerable variability.

## Data Availability

Raw data for these experiments, along with code to access them, have been deposited in a Figshare repository: __________________________________.

## Author Contributions

E.T. performed all surgeries and oversaw animal husbandry. S.A.M. and M.D. contributed to manuscript preparation and data interpretation. D.L.R. devised the experiments, wrote the visual stimulus, collected and analyzed the data, prepared the data for the repository, and wrote the first version of the manuscript.

## Acknowledgements

We thank Matteo Mariani and Alex Huk and for comments on an earlier version of this manuscript.

